# Machine learning based imputation techniques for estimating phylogenetic trees from incomplete distance matrices

**DOI:** 10.1101/744789

**Authors:** Ananya Bhattacharjee, Md. Shamsuzzoha Bayzid

## Abstract

**Background:** Due to the recent advances in sequencing technologies and species tree estimation methods capable of taking gene tree discordance into account, notable progress has been achieved in constructing large scale phylogenetic trees from genome wide data. However, substantial challenges remain in leveraging this huge amount of molecular data. One of the foremost among these challenges is the need for efficient tools that can handle missing data. Popular distance-based methods such as neighbor joining and UPGMA require that the input distance matrix does not contain any missing values.

**Results:** We introduce two highly accurate machine learning based distance imputation techniques. One of our approaches is based on matrix factorization, and the other one is an *autoencoder* based deep learning technique. We evaluate these two techniques on a collection of simulated and biological datasets, and show that our techniques match or improve upon the best alternate techniques for distance imputation. Moreover, our proposed techniques can handle substantial amount of missing data, to the extent where the best alternate methods fail.

**Conclusions:** This study shows for the first time the power and feasibility of applying deep learning techniques for imputing distance matrices. The autoencoder based deep learning technique is highly accurate and scalable to large dataset. We have made these techniques freely available as a cross-platform software (available at https://github.com/Ananya-Bhattacharjee/ImputeDistances).

## Background

Phylogenetic trees, also known as evolutionary trees, represent the evolutionary history of a group of entities (i.e., species, genes, etc.). Phylogenetic trees provide insights into basic biology, including how life evolved, the mechanisms of evolution and how it modifies function and structure etc. One of the ambitious goals of modern science is to construct the “Tree of Life” – the relationships of all organisms on earth. Central to assembling this tree of life is the ability to efficiently analyze the vast amount of genomic data available these days due to the rapid growth rate of newly sequenced genomes.

The field of phylogenetics has experienced tremendous advancements over the last few decades in terms of estimating *gene trees* and *species trees*. Sophisticated and highly accurate statistical methods for reconstructing phylogenetic trees mostly depend on probabilistic models of sequence evolution, and estimate trees using maximum likelihood or Markov Chain Monte Carlo (MCMC) methods (see [1] for example). Various coalescent-based species tree methods with statistical guarantees of returning the true tree with high probability (as the number of genes increases) have been developed, and are increasingly popular [2–11]. However, these methods are not scalable enough to be used with phylogenomic datasets that contain hundreds or thousands of genes and taxa [12,13]. Therefore, developing fast and less computationally demanding, yet reasonably accurate methods remains as one of the foremost challenges in large-scale phylogenomic analyses. Distance-based methods represent an attractive class of methods for large-scale analyses due to their computational efficiency and ease of use. Several studies [10, 11,14–18] have provided support for the considerably good accuracy of distance-based methods, although these methods are generally not as accurate as the computationally demanding Bayesian or likelihood based methods. Distance-based methods can provide reasonably good trees to be used as *guide trees* (also known as *starting trees*) for other sophisticated methods as well as for divide-and-conquer based boosting methods [13, 19–23]. Moreover, under various challenging model conditions, distance-based methods become the only viable option for constructing phylogenetic trees. Whole genome sequences are one such case where traditional approach of multiple sequence alignments may not work [24]. Auch *et al.* [25] proposed a distance-based method to infer phylogeny from whole genome sequences and discussed the potential risks associated with other approaches. Gao *et al.* [26] also introduced a composite vector approach for whole genome data where distances are computed based on the sharing of oligopeptides.

For various practical reasons as discussed above, distance based method has been one of the most popular and widely used techniques, and notable progress have been made in this particular area of phylogenetics [1, 15, 16, 18, 27–29]. Recent works like [30] have made substantial progress towards attaining better accuracy, at least for particular datasets. These improved methods can also be used to obtain information from large-scale single nucleotide polymorphism (SNP) dataset [31].

Missing data is considered as one of the biggest challenges in phylogenomics [32–34]. Missing data can arise from a combination of reasons including data generation protocols, failure of an experimental assay, approaches to taxon and gene sampling, and gene birth and loss [31,35]. The presence of taxa comprising substantial amount of missing (unknown) nucleotides may significantly deteriorate the accuracy of the phylogenetic analysis [34, 36, 37], and can affect branch length estimations in traditional Bayesian methods [38]. Sometimes, presence of missing data can impact the whole arena of phylogenetics, as many studies willingly avoid working with missing data and simply conduct experiments on the available complete dataset [33]. Several paleontology-oriented studies have reported that incomplete taxa can frequently result in poorly resolved phylogenetic relationships [39, 40], and reduces the chance to rebuild the true phylogenetic tree [36].

Several widely-used distance-based methods, including Neighbor Joining [15], UPGMA [27], and BioNJ [16] cannot handle missing data since they require that the distance matrices do not contain and missing entries. However, only a few studies have addressed the imputation of distance values [31, 41]. These works mainly rely on two approaches - direct and indirect. Direct approaches are used by those which try to construct a tree directly from a partially filled distance matrix [1,42]. Indirect approaches, on the other hand, estimate the missing cells at first and then construct phylogenetic tree based on the complete matrix [43, 44]. Some studies like [37] have tried to combine the advantages of both approaches. Recent work like LASSO [31], which uses an algorithm similar to UPGMA, tries to exploit the redundancy in a distance matrix. This method, requiring the assumption of a molecular clock, has been shown to be relatively less accurate by Xia *et al.* [41], as significant differences were observed between the original trees and the trees reconstructed by LASSO from incomplete distance matrices. Xia *et al.* [41] proposed a least square method with multivariate optimization which was shown to achieve high accuracy, even when 10% of the total entries in a distance matrix are missing. However, although this method does not require any molecular clock, it can not determine missing distances when there are sister species with missing distances. Moreover, as we will show in this study, this method is not suitable for distance matrices with substantial amount of missing entries.

In this paper, we propose two statistical and machine learning based approaches to impute missing entries in distant matrices, which do not require any particular assumptions (e.g., molecular clock) and can handle large numbers of missing values. Our techniques are based on *matrix factorization* [45] and *autoencoder* (an unsupervised artificial neural network to learn the underlying representation (encoding) of data) [46]. We report, on an extensive evaluation study using a collection of real biological and simulated dataset, the performance of our methods in comparison with the method proposed by Xia *et al.* [41] (implemented in the DAMBE software package [47,48]) and Kettleborough *et al.* [31] (implemented in the LASSO software package [49]). Experimental results suggest that our methods are more accurate and robust than DAMBE and LASSO under most of the model conditions, and can handle significantly more amount of missing values. This is the only known study that has adapted and leveraged the power of machine learning and deep learning framework in imputing missing values in the context of phylogenetic analyses, and offers the ability to handle large amount of missing values.

## Methods

### Matrix Factorization (MF)

Matrix factorization (MF) has become popular since 2006, when one group of competitors for Netflix Prize that year used the technique [45,50]. Usually being applied in recommender systems [51], this method is used to discover latent features between two interacting entities. Matrix factorization is actually a class of collaborative filtering algorithms [52], which predict users’ future interest by analyzing their past behavior.

Intuitively, there should be some latent features behind how a certain user rates an item. For example, movie ratings by users generally rely on many features including genre, actors, etc. If a certain individual gives high ratings to action movies, we can expect him to do the same to another action movie not rated by him already. Discovering the latent features will thus help predict users’ future preferences.

We adapt this idea to our problem of missing entries in distance matrices. If the distance between two taxa *A* and *B* is not known, we can predict the distance by analyzing their distances with other taxa using the concept of matrix factorization (with appropriate customization).

Let *S* be a set of *N* OTUs (Operational Taxonomic Units). Let, *R* be an |*N*| × |*N*| distance matrix comprising the distances between any two OTUs. If we want to find *K* latent features of distances, we need to find two matrices *X* and *Y*, where the dimensions of *X* and *Y* are |*N*| × *K*. The product of *X* and *Y*^T^ will then approximate *R* as follows.

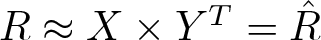

However, as matrix *R* (and *R̂*) has the property where *r*_*ij*_ = *r*_*ji*_ (and *r̂*_*ij*_ = *r̂*_*ji*_), we only consider the lower triangular portion of the matrix. We impute the distance *r̂*_*ij*_ between two OTUs as follows.

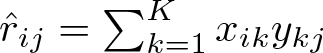

We initialize *X* and *Y* with some random values and try to determine the error between *R* and the product of *P* and *Q*. Then we update those matrices accordingly. We considered squared error as the errors can be both positive and negative. We also add a regularization parameter *β* to avoid overfitting. Thus, we calculate the error as follows.

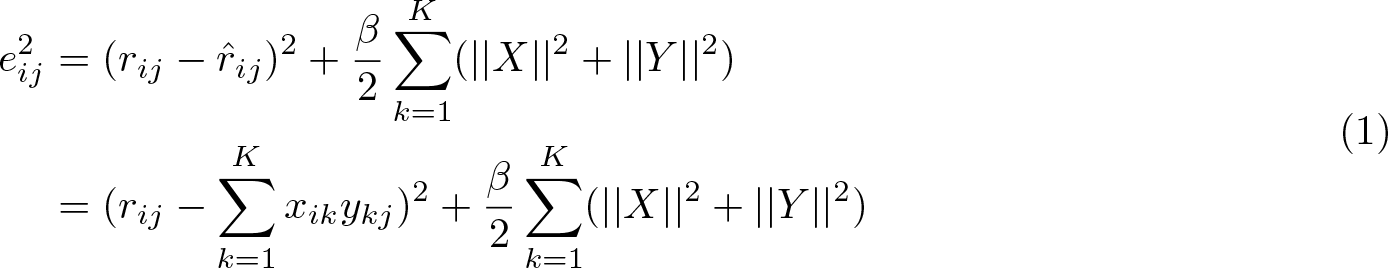

We then obtain the gradient at current values by differentiating Eqn. 1 with respect to *x*_*ik*_ and *y*_*kj*_ separately. We use the following update rules.

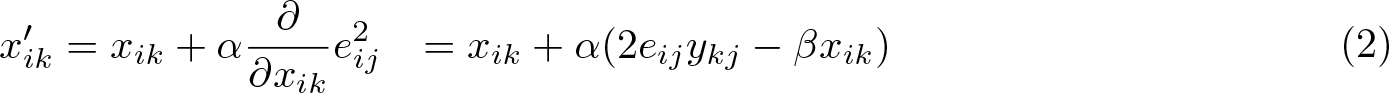

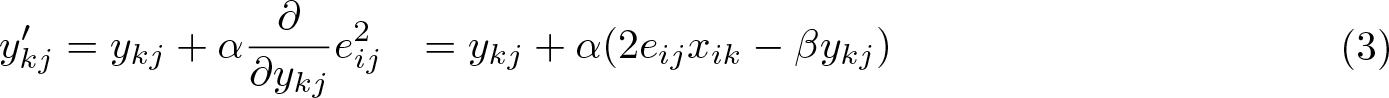

In Equations 2 and 3, *α* is a constant which determines the rate to approach minimum error. We perform the above steps iteratively until the total error *E* (= ∑*e*_*ij*_) converges to a pre-specified threshold value (10^−6^) or 10,000 iterations take place.

Matrix Factorization has previously been used in imputing missing data in various domains of bioinformatics, including analyzing scRNA-seq with missing data [53], handling missing data in genome-wide association studies (GWAS) [54], and identifying cancerous genes [55]. In this study, we successfully adapted this idea for imputing missing entries in a distance matrix for phylogenetic estimation.

### Autoencoder (AE)

Autoencoder (AE) is a type of artificial neural network that learns to copy its input to its output. This is achieved by learning efficient data codings in an unsupervised manner to recreate the input. An autoencoder first compresses the input into a latent space representation and then reconstructs the output from that representation. It tries to learn a function *g*(*f* (*x*)) ≈ *x*, where *f* (*x*) encodes the input and *g*(*f* (*x*)) reconstructs the input using decoder. Figure 1 shows a general overview of autoencoders.

**Figure 1:**
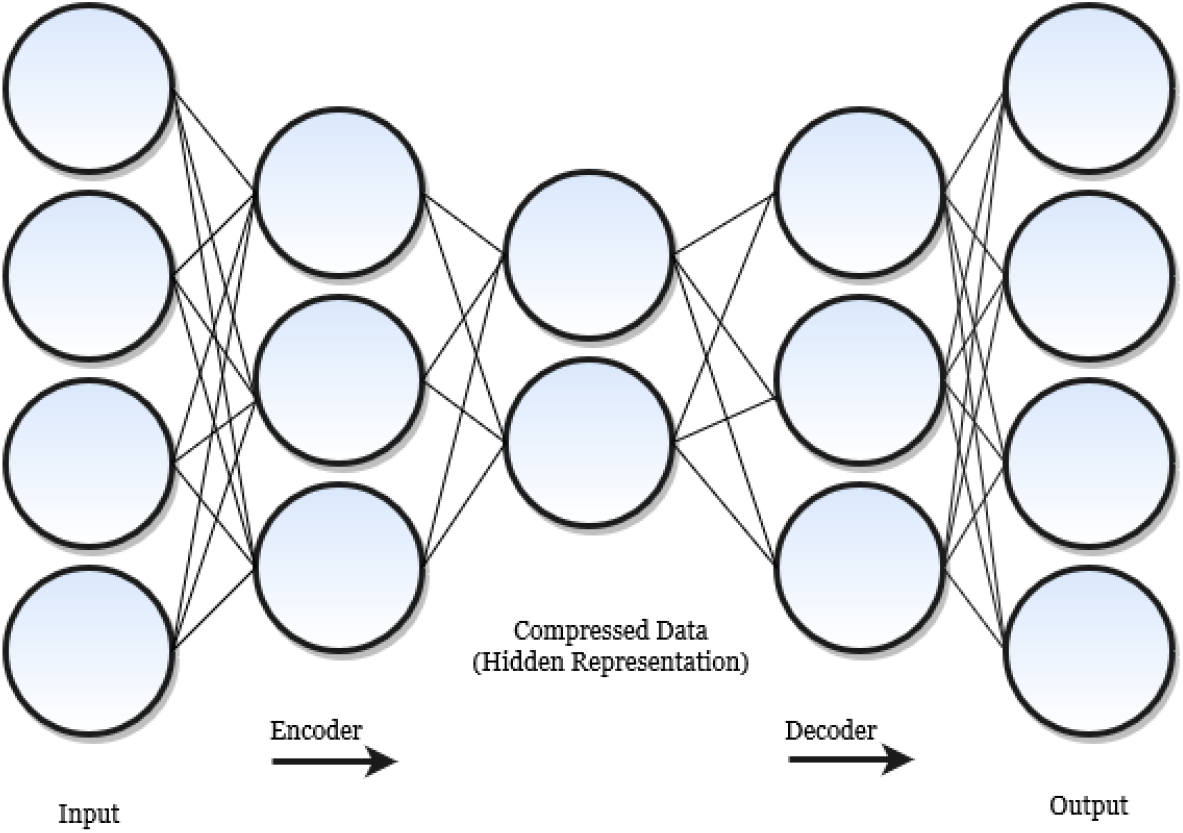
General overview of an autoencoder.

Autoencoder has found various uses in integrative analysis of biomedical big data. Its property of having the ability to reduce dimension and extract non-linear features[56] have been leveraged by many studies. In one oncology study, autoencoders have been able to extract cellular features, which can correlate with drug sensitivity involved with cancer cell lines [57]. Autoencoder was also used to discover two liver cancer sub-types that had distinguishable chances of survival [58]. Moreover, some recent successful data imputation methods have been developed based on autoencoders [59–61]. Autoimpute [59] can be an example which imputes single cell RNA-seq gene expression. Autoencoder-based methods such as [60] and [61] have surpassed older machine learning techniques on various real life datasets.

In this study, we developed an *undercomplete* autoencoder [46] to predict the missing values in distance matrix. The goal of an underdeveloped autoencoder is to learn the most salient features of data by putting a constraint on the amount of information that can flow through the network. There is no need for regularization because they perform maximization of the probability of the data and does not involve copying of the input to the output.

Our architecture has been inspired by an open source library, *FancyImpute* [62]. The model has 3 hidden layers with ReLU (Rectified Linear Unit) *activation functions* [63]. The *dropout rate* is set to 0.75, which appears to work better than other values. Sigmoid function [64] is used as the activation function for output layer. The usual mean squared error (MSE) has been considered for the reconstruction error function. Other than using the predictions provided by the neural network, we have used some predefined weight to update the missing values. A schematic of our model is shown in Fig. 2.

**Figure 2:**
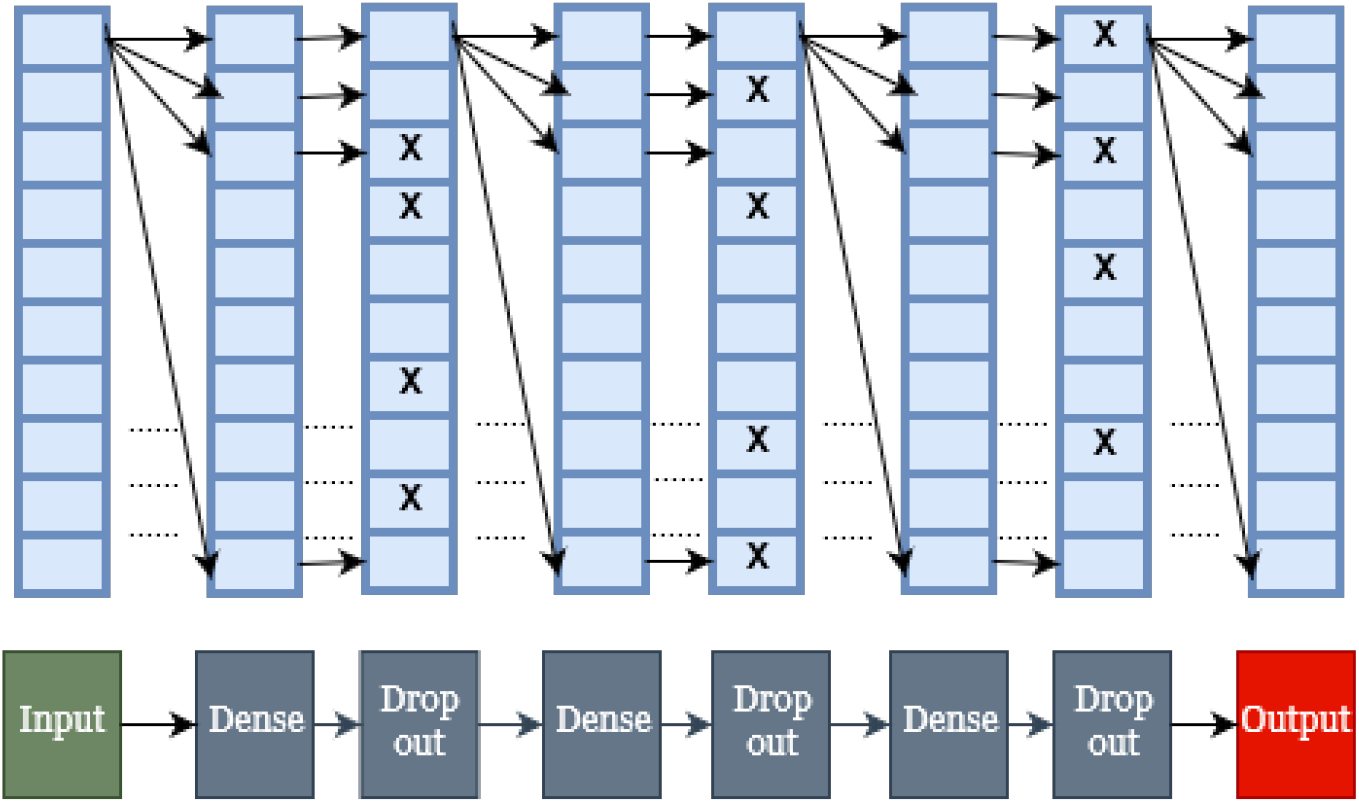
A schematic of our proposed autoencoder model. The *X*’s in the dropout layers symbolically denote that their weights will be set to zero.

## Results

We compared our methods with two of the most accurate alternate methods: 1) the imputation method proposed by Xia *et al.* which is implemented in the software package DAMBE [47, 48], 2) the method proposed by Kettleborough *et al.* [31], which is implemented in the LASSO software package [49]. In this paper, we refer by DAMBE the imputation method proposed by Xia *et al.* [41].

We used a collection of previously studied simulated and biological datasets to evaluate the performance of these methods. We compared the estimated species trees to the model species tree (for the simulated datasets) or to the trees estimated on the full data without any missing entries (for the biological datasets), to evaluate the accuracy of various imputation techniques. We have used normalized Robinson-Foulds (RF) distance [65] to measure the tree error. The RF distance between two trees is the sum of the bipartitions (splits) induced by one tree but not by the other, and vice versa. Normalized RF distance (RF rate) is obtained by dividing the RF distance by the maximum possible RF distance.

Similar to previous studies [41], we generated missing entries in two ways: i) modifying the input sequences in a way that results into missing entries in the distance matrix, and ii) directly deleting entries from a given distance matrix. In a complete distance matrix of *n* taxa, there are 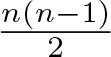 distances since the distance between two entities is symmetric. Similar to previous studies [31, 41], we randomly remove some entries to create partial distance matrices. We can create partial distances by modifying sequence data as well [41] (see the next sub-section on datasets). We have used FastME [18, 29] to construct trees from complete distance matrices.

### Datasets

We have used a set of mitochondrial COI and CytB sequences from 10 Hawaiian katydid species in the genus *Banza* along with four outgroup species. This dataset, comprising 24 operational taxonomic units (OTUs) and 10 genes which evolved under the HKY85 model [66], was previously used in [41]. In order to evaluate the relative performance, we followed exactly the same process used by Xia *et al.* [41] for modifying the sequences to create missing entries in distance matrices. However, Xia *et al.* only generated 30 missing entries in the matrix, whereas we analyzed a wide range of missing entries (10 ∼ 140).

We now explain how missing values were introduced by modifying the sequences both in [41] and this study. A set of mitochondrial COI and CytB sequences was used for the complete dataset of 24 OTUs. If we remove COI sequence from a taxon A and CytB sequence from another taxon B, then (A, B) pair does not share any homologous sites which results into a missing entry in the corresponding distance matrix. Thus, if we remove COI sequence from *n*_1_ taxa and remove CytB sequence from a different set of *n*_2_ taxa, we will have *n*_1_ × *n*_2_ missing entries in the distance matrix. From the sequences, we created incomplete matrices based on the MLCompositeTN93 (TN93) model [67]. TN93 model holds the assumption of a complex but specific model of nucleotide substitution. The distance formula is derived under the homogeneity assumption, which means that the pattern of nucleotide substitution has not changed in the evolutionary history of the observed sequences [68, 69]. We used *MEGA-X* [69–71] to introduce missing entries in the distance matrices.

We used another set of simulated dataset based on a biological dataset (37-taxon mammalian dataset [72]) that was generated and subsequently analyzed in prior studies [9, 13, 73, 74]. This dataset was generated under the multi-species coalescent model [75] with various model conditions reflecting varying amounts of gene tree discordance resulting from the incomplete lineage sorting (ILS) [76]. This collection of dataset was simulated by taking the species tree estimated by MP-EST [7] on the biological dataset studied in Song *et al.* [72]. This species tree had branch lengths in coalescent units, that were scaled (multiplying or dividing by two) to vary the amount of ILS (shorter branch lengths result into more ILS). The basic model condition with moderate amount of ILS is referred to as 1X and the model conditions with higher and lower amounts of ILS are denoted by 0.5X and 2X, respectively. For each model condition, we used 10 replicates of data each containing 37 sequences.

In addition to the TN93 model used in previous studies [41], we also applied the LogDet method [77] to observe how they affect the imputation process. LogDet is often considered superior as it does not associate itself with the assumptions held by TN93, although it may overestimate distances in certain cases [68]. We analyzed a range of missing entries: 36 (6 × 6), 100 (10 × 10), 225 (15 × 15), and 342 (19 × 18). We also analyzed three distance matrices which were computed from aligned sequences (from Carnivores, Baculovirus, and mtDNAPri3F84SE) and were used in previous studies [78, 79]. The numbers of taxa in these matrices ranges from 7 to 10.

Various numbers of distance values were randomly removed to introduce missing data.

### Results on Sequence Input

Table 1 shows the results on 24 OTUs for a wide range of missing entries (10 ∼ 140). For this particular dataset, MF showed superior performance on small to moderate numbers of missing entries (0 ∼ 40), LASSO matched or improved upon the other methods for moderate to high numbers of missing entriess (50 ∼ 110), and AE outperformed others in the presence of higher amounts of missing data (110 ∼ 140).

**Table 1:** RF rates of different methods on the 24 taxa dataset with varying numbers of missing entries. The best RF rates for various model conditions are shown in boldface.

| #Taxa | #Entries | #Missing Entries | RF Rate |  |  |  |
| --- | --- | --- | --- | --- | --- | --- |
|  |  |  | DAMBE | LASSO | MF | AE |
| 24 | 276 | 10 | 0.0476 | 0.2857 | **0** | 0.0952 |
|  |  | 20 | **0.1429** | 0.3333 | 0.1905 | 0.2381 |
|  |  | 30 | 0.2381 | 0.3333 | **0.1905** | 0.2381 |
|  |  | 40 | **0.2857** | 0.3333 | **0.2857** | 0.3333 |
|  |  | 50 | 0.3333 | **0.1905** | 0.4286 | 0.3333 |
|  |  | 60 | 0.2857 | **0.2381** | 0.3333 | 0.381 |
|  |  | 70 | 0.4286 | **0.2857** | 0.5714 | 0.381 |
|  |  | 80 | 0.4762 | **0.381** | 0.6667 | **0.381** |
|  |  | 90 | 0.5714 | **0.5238** | **0.5238** | **0.5238** |
|  |  | 100 | **0.5714** | **0.5714** | 0.7143 | 0.6190 |
|  |  | 110 | 0.7143 | **0.6190** | 0.8571 | **0.6190** |
|  |  | 120 | 0.8095 | 0.7619 | 0.8571 | **0.7143** |
|  |  | 130 | 0.8571 | **0.7619** | 0.8095 | **0.7619** |
|  |  | 140 | N/A | N/A | **0.7619** | **0.7619** |

For 30 missing entries (which was the case analyzed in [41]), MF recovered 81% of the true bipartitions, whereas DAMBE and LASSO recovered 76% and 67% bipartitions respectively. Fig. 3 illustrates the differences among the trees constructed by various methods with 30 missing values. MF estimated tree is more closer to the tree estimated on the full dataset compared to DAMBE and AE up to 40 missing entries. Notably, with 10 missing entries, MF was able to reconstruct the correct tree whereas DAMBE and AE incurred 5% and 10% errors, respectively. However, as we increase the number of missing entries, DAMBE started to outperform MF, and AE started to outperform both DAMBE and MF. Moreover, for moderate and high numbers of missing taxa (50 ∼ 110), LASSO showed the best performance in recovering true bipartitions, although sometimes MF and AE were equally good. When one-third of the entries in the distance matrix are missing, LASSO, MF, and AE recovered almost 48% bipartitions, whereas DAMBE recovered 43% bipartitions. Another important point is that DAMBE can not impute distances when more than 50% of the total entries are missing. LASSO’s performance is not promising in this case either, because LASSO can not construct a tree on the full set of taxa, resulting in an incomplete tree. Therefore, we could not consider the trees produced by LASSO when more than 50% of the entries are missing. On the other hand, both MF and AE were able to reconstruct around 25% of the true bipartitions even when more than 50% of the entries are missing. Although, more than 50% missing entries in a distance matrix may not be a very common model condition, the ability to handle arbitrarily large amounts of missing data advances the state-of-the-art in imputation techniques. The trees generated by MF and AE on dataset with higher amounts of missing data can be used as starting trees for further improvements.

**Figure 3:**
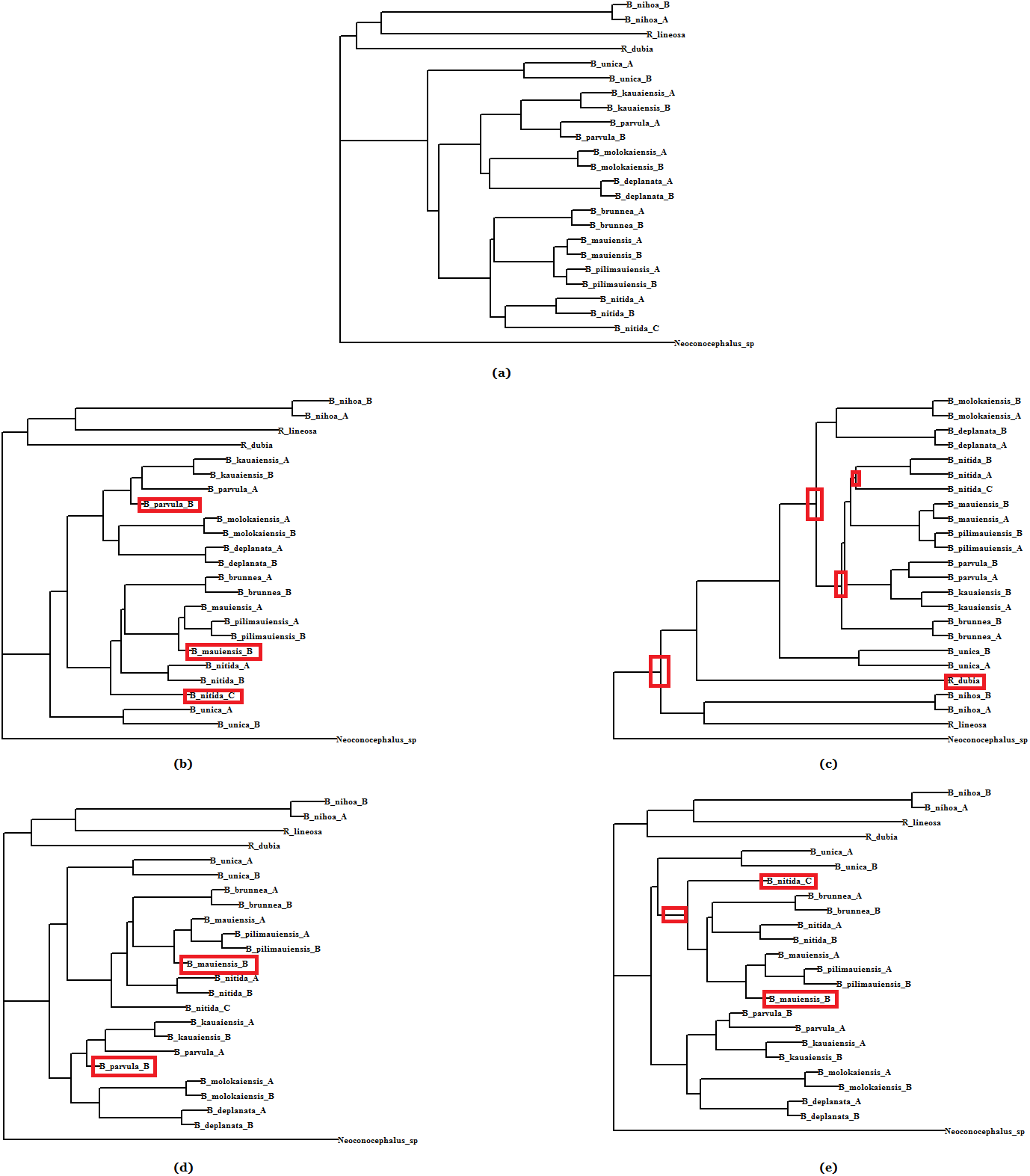
Phylogenetic trees estimated on the full and incomplete dataset (30 missing entries) with 24 OTUs from 10 Hawaiian katydid species. (a) Tree estimated from the full data (complete distance matrix), (b) - (e) trees reconstructed from incomplete distance matrix by DAMBE, LASSO, MF, and AE, respectively. Red rectangles highlight the inconsistencies with the tree on the full dataset.

Results on 37-taxon simulated dataset with varying amounts of ILS, two different evolution models and varying numbers of missing entries, are demonstrated in Tables 2, 3, and 4. MF and AE are comparable or better than DAMBE in most of the case. One noticeable aspect in these tables is that, unlike the 24 OTUs dataset, LASSO performed very poorly in all cases, providing the worst recovery rate of true bipartitions. As DAMBE and LASSO can not handle distance matrices with more than 50% missing entries, only MF and AE were able to run on the distance matrices with 342 missing entries, albeit the RF rates are very high (due to the lack of sufficient phylogenetic information present in the highly incomplete distance matrix). MF could not recover any internal branches on the 1X dataset with 50% missing entries. AE, on the other hand, was able to reconstruct around 15% bipartitions. Another observation, within the scope of the experiments performed in this study, is that the amounts of ILS do not have any significant impact on the performance of these imputation techniques. However, more experiments and analyses are required to further investigate the impact of ILS on the tree estimation from incomplete distance matrices.

**Table 2:**
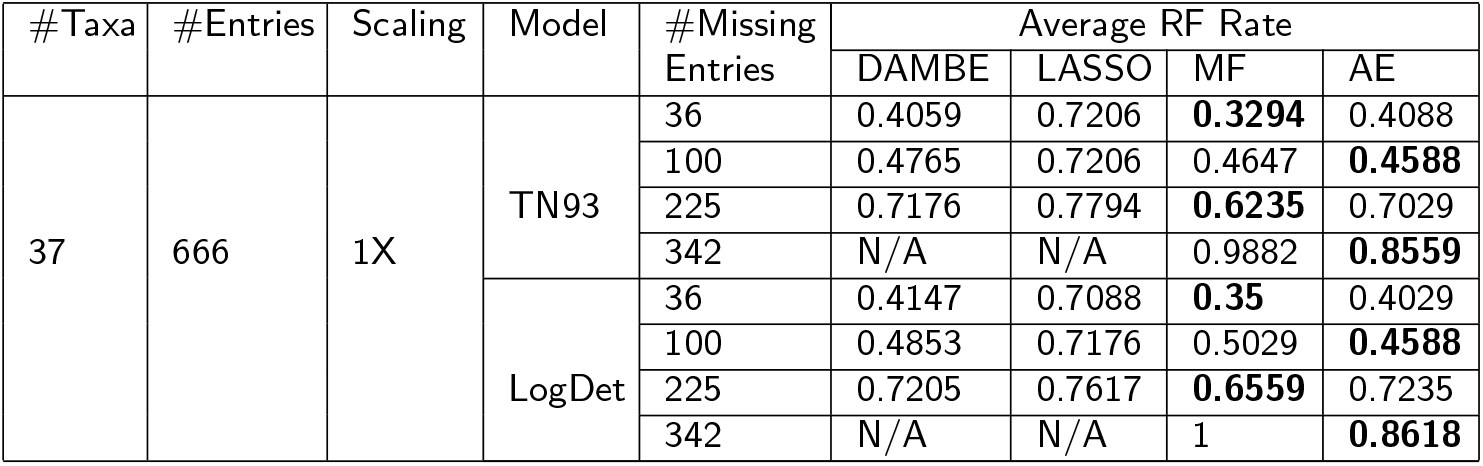
Average RF rates of different methods on the 37-taxon, 1X dataset (moderate amount of ILS) for varying numbers of missing entries and two different sequence evolution models. We show the average RF rates over 10 replicates. The best RF rates for various model conditions are shown in boldface.

| #Taxa | #Entries | Scaling | Model | #Missing Entries | Average RF Rate |  |  |  |
| --- | --- | --- | --- | --- | --- | --- | --- | --- |
|  |  |  |  |  | DAMBE | LASSO | MF | AE |
| 37 | 666 | 1X | TN93 | 36 | 0.4059 | 0.7206 | **0.3294** | 0.4088 |
|  |  |  |  | 100 | 0.4765 | 0.7206 | 0.4647 | **0.4588** |
|  |  |  |  | 225 | 0.7176 | 0.7794 | **0.6235** | 0.7029 |
|  |  |  |  | 342 | N/A | N/A | 0.9882 | **0.8559** |
|  |  |  | LogDet | 36 | 0.4147 | 0.7088 | **0.35** | 0.4029 |
|  |  |  |  | 100 | 0.4853 | 0.7176 | 0.5029 | **0.4588** |
|  |  |  |  | 225 | 0.7205 | 0.7617 | **0.6559** | 0.7235 |
|  |  |  |  | 342 | N/A | N/A | 1 | **0.8618** |

**Table 3:** Average RF rates of different methods on the 37-taxon, 0.5X dataset (high ILS) for varying numbers of missing entries and two different sequence evolution models. We show the average RF rates over 10 replicates. The best RF rates for various model conditions are shown in boldface.

| #Taxa | #Entries | Scaling | Model | #Missing Entries | Average RF Rate |  |  |  |
| --- | --- | --- | --- | --- | --- | --- | --- | --- |
|  |  |  |  |  | DAMBE | LASSO | MF | AE |
| 37 | 666 | 0.5X | TN93 | 36 | 0.4529 | 0.6941 | **0.3471** | 0.4294 |
|  |  |  |  | 100 | **0.4912** | 0.7206 | 0.5029 | 0.5353 |
|  |  |  |  | 225 | 0.6618 | 0.7588 | **0.6235** | 0.7147 |
|  |  |  |  | 342 | N/A | N/A | 0.9971 | **0.8412** |
|  |  |  | LogDet | 36 | 0.45 | 0.6824 | **0.35** | 0.4176 |
|  |  |  |  | 100 | **0.4853** | 0.7059 | 0.5236 | 0.5147 |
|  |  |  |  | 225 | **0.6441** | 0.7588 | 0.6559 | 0.7029 |
|  |  |  |  | 342 | N/A | N/A | 0.9853 | **0.8412** |

**Table 4:**
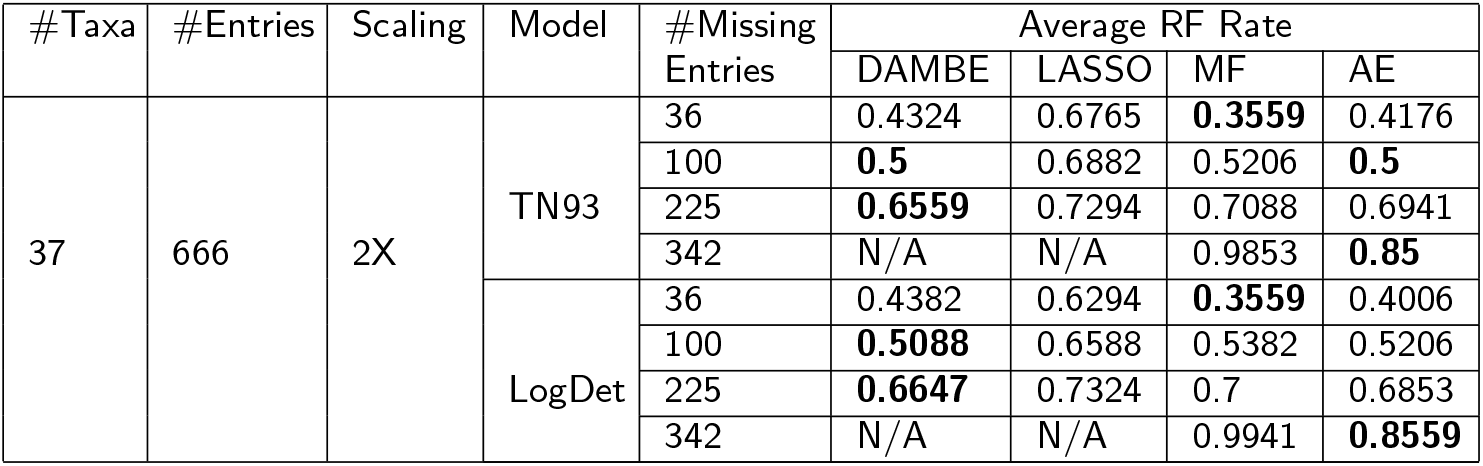
Average RF rates of different methods on the 37-taxon, 2X dataset (low ILS) for varying numbers of missing entries and two different sequence evolution models. We show the average RF rates over 10 replicates. The best RF rates for various model conditions are shown in boldface.

| #Taxa | #Entries | Scaling | Model | #Missing Entries | Average RF Rate |  |  |  |
| --- | --- | --- | --- | --- | --- | --- | --- | --- |
|  |  |  |  |  | DAMBE | LASSO | MF | AE |
| 37 | 666 | 2X | TN93 | 36 | 0.4324 | 0.6765 | **0.3559** | 0.4176 |
|  |  |  |  | 100 | **0.5** | 0.6882 | 0.5206 | **0.5** |
|  |  |  |  | 225 | **0.6559** | 0.7294 | 0.7088 | 0.6941 |
|  |  |  |  | 342 | N/A | N/A | 0.9853 | **0.85** |
|  |  |  | LogDet | 36 | 0.4382 | 0.6294 | **0.3559** | 0.4006 |
|  |  |  |  | 100 | **0.5088** | 0.6588 | 0.5382 | 0.5206 |
|  |  |  |  | 225 | **0.6647** | 0.7324 | 0.7 | 0.6853 |
|  |  |  |  | 342 | N/A | N/A | 0.9941 | **0.8559** |

We also analyzed the impact of two widely used sequence evolution models (TN93 and LogDet) on the performance of the proposed imputation techniques. MF performed poorly on LogDet model compared to the TN93 model, as 17 out of 24 cases have LogDet producing a greater RF rate than TN93. AE, on the other hand, shows similar (on 1X model) or slightly better (on 0.5X and 2X models) performance under LogDet construction. DAMBE achieved performance improvement under LogDet construction only for the 0.5X model condition (Table 3) and the opposite trend is observed for the 1X and 2X model conditions (Table 2 and 4), albeit the differences are very small. Although LASSO performed poorly on these dataset, LogDet model helped it achieve a slightly better performance than TN93.

### Results on Distance Matrix Input

We analyzed three separate distance matrices which were computed from the gene sequences from Carnivores, Baculovirus and mtDNAPri3F84SE, and were analyzed in previous studies [41]. We show the results in Tables 5, 6, and 7. For the carnivores dataset (Table 5), LASSO and AE produced the best results except for one case with 15 missing entries where DAMBE was better than others. Even with more than 50% missing entries, AE was able to reconstruct 30% of the true bipartitions. The performance of MF was worse than LASSO, AE, and DAMBE on this particular dataset. On the Baculovirus dataset, DAMBE achieved the lowest RF rates for relatively lower numbers of missing entries. However, as we increase the amount of missing entries, MF and AE starts to outperform other methods. On the mtDNAPri3F84SE dataset, the performances of these methods were mixed, and no method consistently outperformed the others. However, DAMBE and LASSO achieved better performance than MF and AE. Notably, AE was able to reconstruct 35% and 50% true bipartitions on Baculovirus and mtDNAPri3F84SE dataset even when more than 50% of the distance values were missing.

**Table 5:** RF rates of different methods on the Carnivores dataset. The best RF rates for various model conditions are shown in boldface.

| #Taxa | #Entries | #Missing Entries | RF Rate |  |  |  |
| --- | --- | --- | --- | --- | --- | --- |
|  |  |  | DAMBE | LASSO | MF | AE |
| 10 | 45 | 5 | 0.5714 | **0.1429** | 0.4286 | 0.4286 |
|  |  | 10 | 0.4286 | **0.1429** | 0.7143 | **0.1429** |
|  |  | 15 | **0** | 0.2857 | 0.7143 | 0.5714 |
|  |  | 20 | 0.8571 | **0.1429** | 0.5714 | 0.5714 |
|  |  | 25 | N/A | N/A | 0.8571 | **0.7143** |

**Table 6:** RF rates of different methods on the Baculovirus dataset. The best RF rates for various model conditions are shown in boldface.

| #Taxa | #Entries | #Missing Entries | RF Rate |  |  |  |
| --- | --- | --- | --- | --- | --- | --- |
|  |  |  | DAMBE | LASSO | MF | AE |
| 9 | 36 | 4 | **0** | 0.1667 | 0.1667 | 0.1667 |
|  |  | 8 | **0.1667** | **0.1667** | 0.5 | 0.3333 |
|  |  | 12 | 0.6667 | 0.6667 | **0.5** | 0.6667 |
|  |  | 16 | 0.8333 | 0.6667 | 0.6667 | **0.5** |
|  |  | 20 | N/A | N/A | 0.8333 | **0.6667** |

**Table 7:** RF rates of different methods on the mtDNAPri3F84SE dataset. The best RF rates for various model conditions are shown in boldface.

| #Taxa | #Entries | #Missing Entries | RF Rate |  |  |  |
| --- | --- | --- | --- | --- | --- | --- |
|  |  |  | DAMBE | LASSO | MF | AE |
| 7 | 21 | 2 | **0** | **0** | **0** | **0** |
|  |  | 5 | 0.5 | **0.25** | 0.5 | 0.75 |
|  |  | 7 | **0** | 0.25 | 0.75 | 0.25 |
|  |  | 10 | 0.75 | **0.25** | 0.75 | 0.75 |
|  |  | 12 | N/A | N/A | 0.75 | **0.5** |

### Running Time

We performed the experiments on a computer with i5-3230M, 2.6 GHz CPU with 12 GB RAM. Among these four methods, MF was the slowest method. The running time of MF on the 24-taxon dataset ranges between 7 ∼ 15 minutes for various numbers of missing entries. DAMBE takes only a few seconds with 10 missing entries, but as we increase the number of missing entries to 130, the running time of DAMBE increases to 2 minutes. AE was faster, requiring only around 30 seconds for this dataset. LASSO was the fastest, taking only a second. Notably, unlike MF and DAMBE, the running times of AE and LASSO do not change as we increase the number of missing entries.

For the 37-taxon dataset, MF takes around 30 minutes while DAMBE takes 12 ∼ 15 minutes. AE is faster than MF and DAMBE, taking only around 45 seconds. LASSO was the fastest method which took only a second. For the relatively smaller matrices presented in Sec., DAMBE is very fast, and finished in a second. MF took around 45 seconds, and AE took 20 seconds. Overall, the running time of LASSO and AE are better than others and are less sensitive to the numbers of taxa and the numbers of missing entries.

## Discussion

We extensively evaluated our proposed methods on a collection of real and simulated dataset. Previous studies like [41] and [31] have limited their evaluation studies to only one or two datasets with limited numbers of taxa. Moreover, previous studies also limit their results to 10% missing entries. We tried to address these issues by testing our methods as well as the previous ones on five different datasets with different challenging model conditions and applied various missingness mechanisms. We have tested our methods for a wide range of missing entries. We also worked on a 37-taxon mammalian dataset, whereas previous comparative studies were limited to 26 taxa. Furthermore, we analyzed the impact of varying amounts of ILS on the performance of various imputation techniques.

In general, MF and AE seem to be robust across various dataset and model conditions. DAMBE was comparable to MF and AE when the numbers of missing entries were relatively small. However, in general, DAMBE does not perform well with moderate to high numbers of missing entries. Although LASSO was previously shown to be less accurate than DAMBE in [41], we observed mixed performance, and found LASSO performing better than DAMBE in several cases. For lower numbers of taxa, LASSO works very well, even when 25-45% entries are missing. But on the 37-taxon dataset, LASSO consistently performed poorly compared to other methods. Our proposed methods did not show any such indication of obvious poor performance in any particular dataset. Even on the model conditions where LASSO and DAMBE achieved better performance, MF and AE achieved competitive accuracy. MF works especially well when the number of missing entries is small. AE, on the other hand, shows good performance overall, and does particularly well when 50% or more distances are missing.

Another important aspect is both DAMBE and LASSO failed to handle distance matrices with more than 50% missing entries. But our methods have no such limitation. More often than not, sequence data contain substantial amounts of missing information, resulting into distance matrices with lots of missing entries. We understand that, in the presence of a substantial number of missing entries in a distance matrix, researchers will tend to approach the trees with extreme care. However, the ability to construct trees in the presence of arbitrarily large numbers of missing entries will help us estimate starting trees on extremely challenging model conditions with high levels of missing entries. These starting/guide trees can be improved with further analysis (for example, divide-and-conquer based boosting techniques [13, 19–22]).

Although we investigated a collection of dataset under various practical model conditions (more than the number of model conditions analyzed in previous studies), this study can be expanded in several directions. Future works will need to investigate how to help the researchers choose the right imputation approaches for various model conditions. This study investigated relatively long sequences (1500 ∼ 2600 bp); subsequent studies should investigate the relative performance of methods on very short sequences. This study analyzed small to moderate sized dataset (7 ∼ 37 taxa). Larger dataset with hundreds of taxa need to be analyzed, especially to demonstrate the power of machine learning techniques in leveraging the latent features of phylogenetic data. We leave these as future works.

## Conclusions

In this study we have presented two imputation techniques, inspired from matrix factorization and deep learning architecture, to reconstruct phylogenetic trees from partial distance matrices. Experimental results using both simulated and real biological dataset show that our models match or improve upon alternate best techniques under varying model conditions (numbers of taxa, sequence lengths, gene tree discordance, DNA sequence evolution models, etc.), and missingness mechanisms.

Estimating phylogenetic trees in the presence of missing data is sufficiently complex and hence existing methods cannot fully comprehend or predict the relationships among the taxa from partial distance matrices. Thus the goal here should be the creation of an appropriate model to capture the underlying data distribution; the model should account for as much phylogenetic data as possible to impute the missing entries. This view emphasizes the importance of machine learning (ML) for distance matrix imputation. Moreover, we aimed for developing appropriate unsupervised models. Unsupervised learning approaches have advantages over supervised methods particularly when the data are heterogeneous, which are often so with various phylogenetic dataset and therefore the supervised models trained on distance matrices on a particular set of taxa may not be generalizable to impute missing entries in the distance matrices on other set of taxa.

We have shown that MF and AE can handle very high amount of missing data. Unlike other methods [31], our proposed methods do not require the molecular clock assumption. Moreover, deep architecture like autoencoders are able to automatically learn latent representations and complex inter-variable associations, which is not possible using other methods. Our autoencoder based method is also scalable to large datasets. Considering the rapidly increasing growth of phylogenomic dataset, and the prevalence of accompanying missing data, the timing of our proposed approaches seems appropriate. Thus, given the demonstrated potential in tree reconstruction in the face of missing data, we believe that our proposed techniques represent a major step towards solving real world instances in phylogenomics.

## Declarations

### Availability of data and material

The proposed methods is freely available as open source code at https://github.com/Ananya-Bhattacharjee/ImputeDistanceshttps://github.com/Ananya-Bhattacharjee/ImputeDistances. All the dataset analyzed in this paper are from previously published studies and are publicly available.

### Authors’ contributions

MSB and AB conceived the study; AB and MSB designed the methods; AB implemented the methods and performed the experiments; MSB and AB interpreted the results; MSB and AB wrote the paper.

### Ethics approval and consent to participate

Not applicable.

### Consent for publication

Not applicable.

## Acknowledgements

Not applicable.

